# Improving Therapeutic Protein Secretion in the Probiotic Yeast *Saccharomyces boulardii* using a Multifactorial Engineering Approach

**DOI:** 10.1101/2022.12.30.522352

**Authors:** Deniz Durmusoglu, Ibrahim Al’Abri, Taufika Islam Williams, Leonard B. Collins, José L. Martínez, Nathan Crook

**Author notes:** Correspondence, Correspondence during review.

## Abstract

The probiotic yeast *Saccharomyces boulardii* (*Sb*) is a promising chassis to deliver therapeutic proteins to the gut due to *Sb*’s innate therapeutic properties, resistance to phage and antibiotics, and high protein secretion capacity. To maintain therapeutic efficacy in the context of challenges such as washout, low rates of diffusion, weak target binding, and/or high rates of proteolysis, it is desirable to engineer *Sb* strains with enhanced levels of protein secretion. In this work, we explored genetic modifications in both *cis*- (i.e., to the expression cassette of the secreted protein) and *trans*- (i.e., to the *Sb* genome) that enhance *Sb*’s ability to secrete proteins, taking a *Clostridioides difficile* Toxin A neutralizing peptide (NPA) as our model therapeutic. First, by modulating the copy number of the NPA expression cassette, we found NPA concentrations in the supernatant could be varied by 6-fold (76-458 mg/L) in microbioreactor fermentations. In the context of high NPA copy number, we found a previously-developed collection of native and synthetic secretion signals could further tune NPA secretion between 121 - 463 mg/L. Then, guided by prior knowledge of *S. cerevisiae*’s secretion mechanisms, we generated a library of homozygous single gene deletion strains, the most productive of which achieved 2297 mg/L secretory production of NPA. We then expanded on this library by performing combinatorial gene deletions, supplemented by proteomics experiments. We ultimately constructed a quadruple protease-deficient *Sb* strain that produces 5045 mg/L secretory NPA, an improvement of >10-fold over wild-type *Sb*. Overall, this work systematically explores a broad collection of engineering strategies to improve protein secretion in *Sb* and highlights the ability of proteomics to highlight under-explored mediators of this process. In doing so, we created a set of probiotic strains that are capable of delivering a wide range of protein titers and therefore furthers the ability of *Sb* to deliver therapeutics to the gut and other settings to which it is adapted.

## 1. Introduction

Engineered live biotherapeutic products (eLBPs) are living organisms that are genetically engineered to prevent, diagnose, and/or treat disease. In doing so, they can produce therapeutics ranging from small molecules, such as vitamins and antimicrobials, to large molecules such as interleukins and antibodies (Charbonneau et al., 2020; Durmusoglu et al., 2021; Sola-Oladokun et al., 2017). Via *in situ* drug synthesis using organisms that are often cheaper to produce, transport, and store than the drugs themselves, eLBPs promise to reduce side effects and cost for a broad range of health conditions. Currently, the majority of eLBP candidates in clinical trials are bacterial (Cubillos-Ruiz et al., 2021). While bacteria are ideal for many *in situ* applications (e.g. those requiring high numerical abundance, high growth rates, or bacterial metabolic pathways), bacteria have important limitations such as susceptibility to bacteriophage and antibiotics, often limited protein secretion rates, and difficulty in performing post-translational modification of proteins (Brunk et al., 2018).

*Saccharomyces boulardii* (*Sb*) is a probiotic strain of yeast that was isolated from lychee and mangosteen in 1923 by Henri Boulard. While exhibiting 99% genomic similarity to *Saccharomyces cerevisiae* (*Sc*), *Sb* differs substantially from *Sc* phenotypically. *Sb* has a thicker cell wall and better tolerates low pH and body temperature, providing an advantage to survive in the human gut (Fietto et al., 2004; Hudson et al., 2016). *Sb* is available worldwide as a dietary supplement and is often used as an adjunctive therapy for ulcerative colitis, diarrhea and recurrent *Clostridioides difficile* (*C. difficile*) infections (Kabbani et al., 2017; Kelly et al., 2019). Like *Sc*, *Sb* is not susceptible to bacteriophages nor antibiotics and can be programmed to secrete recombinant proteins (Blackburn & Avery, 2003). These features make *Sb* a strong candidate for *in situ* biomanufacturing of therapeutics in the GI tract. Recently, *Sb* has been engineered to secrete human synthetic lysozyme, antimicrobial Leucocin C, and HIV-1 Gag in culture and deliver β-carotene, IL-10, atrial natriuretic peptide, and nanobodies against *C. difficile* toxins in rodent models (K. Chen et al., 2020; Durmusoglu, Al’Abri, et al., 2021; R. Li et al., 2020; J.-J. Liu et al., 2016; Palma et al., 2019). These examples demonstrate the promise of *Sb* as an eLBP chassis, complementary to bacteria, for production and secretion of a range of biomolecules with varying structures in the gut.

The concentration of drug released to the desired site is one of the most important metrics (pharmacokinetics) in developing a drug delivery technology (eLBPs included). Here, we focus on biologic (i.e., protein-based) drugs due to their use in treating gut diseases (Al-Bawardy et al., 2021) and their conserved route for export within an eLBP species. Within the gut, pH, oxygen, and nutrient gradients, as well as an abundance of commensal microbes and proteases, combine to decrease the effective concentration of the secreted protein cargo. In order to mitigate this, eLBPs are often engineered to improve their production and secretion capacities through vector design and/or genome modifications, particularly to the secretory pathway (K. Chen et al., 2020; Geldart et al., 2018; Karlskås et al., 2014; C.-H. Liu et al., 2020). Such approaches have been extensively studied for bacterial eLBPs, but the strategies to improve protein secretion in the eukaryotic eLBP *Sb* have been less extensively explored.

For a protein to be secreted by yeast, four major steps must take place (**Figure 1**) (Hou et al., 2012a; Kastberg et al., 2022). First, the mRNA for the desired protein is synthesized using a genomic or plasmidic template (**Figure 1.1**), with an increased copy number of the DNA template often increasing the amount of mRNA that is synthesized. mRNAs are then transported to the cytoplasm where they are translated into a protein product, that is then translocated into the endoplasmic reticulum (ER) lumen (**Figure 1.2**). This translocation can either occur post-translationally or co-translationally. In post-translational translocation, polypeptides are translocated into the ER lumen after their synthesis is complete (Johnson et al., 2013). In co-translational translocation, insertion of the polypeptide into the ER lumen occurs simultaneously with translation (Nyathi et al., 2013). Which translocation type occurs is determined by the secretion signal sequence at the N-terminus of the recombinant protein (Rapoport et al., 2017). Once in the ER lumen, signal sequence cleavage, glycosylation, disulfide bond formation, and folding occur (Braakman & Hebert, 2013). Quality control (QC) machinery in the ER guides over-accumulated, misfolded, or unfolded proteins to degradation processes such as endoplasmic reticulum associated degradation (ERAD) to ensure ER balance. In particular, recombinant proteins are often recognized as aberrant by QC machinery and may be prematurely directed to ERAD, hindering their trafficking along the secretory pathway. Proteins that are successfully processed in the ER are then trafficked to the Golgi, where they are further glucosylated and mannosylated (**Figure 1.3**). Retrograde trafficking within the Golgi provides additional time for proteins to undergo these modifications, however extensive glycosylation may limit protein secretion (Stanley, 2011). After the Golgi, proteins are either trafficked to the plasma membrane to be secreted, or mis-sorted to the early endosome, which later develops into the vacuole, through the vacuole protein sorting pathway (Bowers & Stevens, 2005) (**Figure 1.4**). Inside the vacuole, proteins encounter a variety of proteases, collectively comprising vacuolar protein degradation (Hecht et al., 2014). Secreted proteins also encounter extracellular proteases and can be degraded outside the cell as well (Sinha et al., 2005). Collectively, this prior knowledge points to “sinks” in the secretory pathway for potential removal in *Sb*.

**Figure 1.**
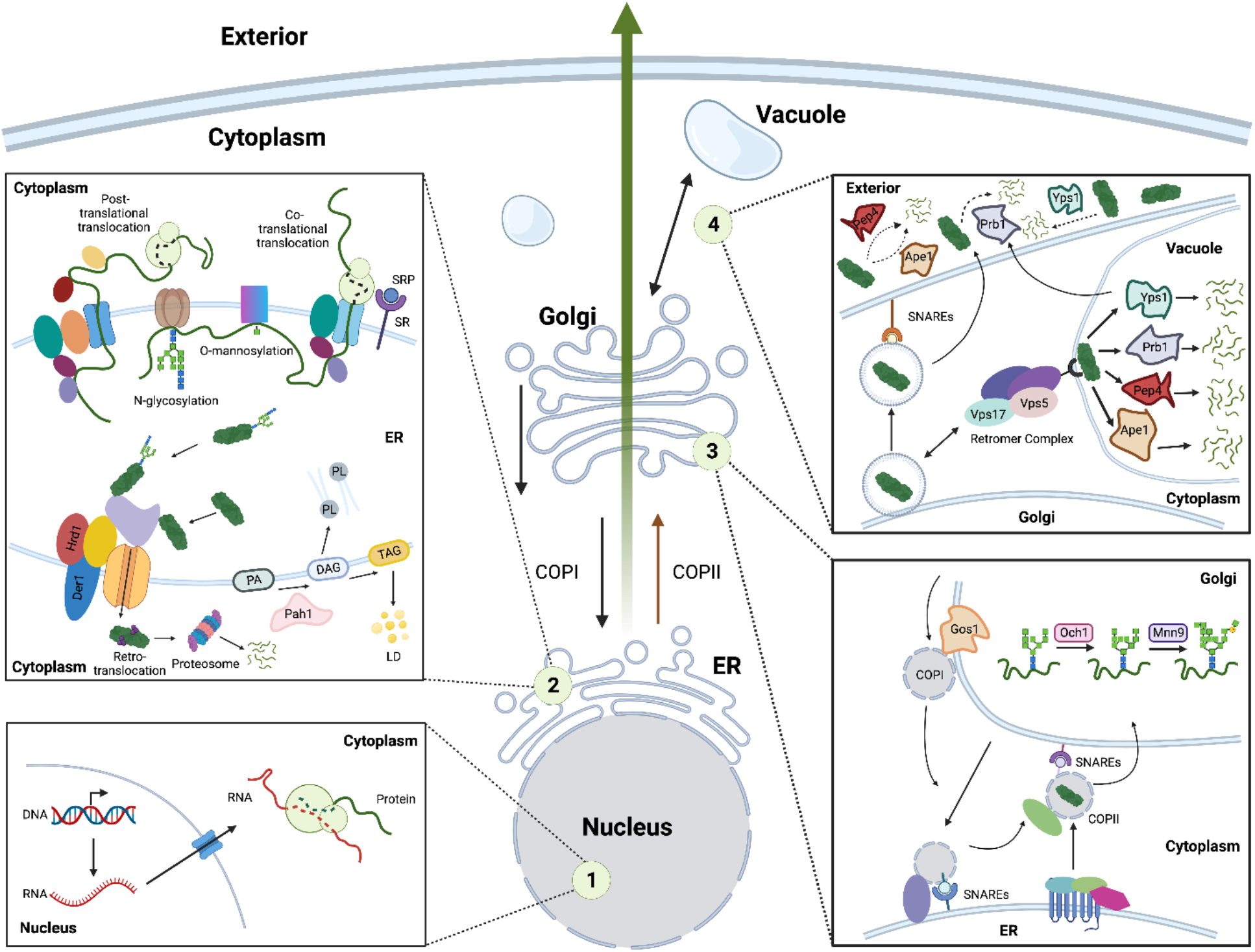
Overview of key steps explored in this study to improve peptide secretion in *Sb*. (1) Transcription of the gene encoding for the secreted protein, located on the genome or on multicopy plasmids. (2) Translocation and modifications that occur in the ER. Nascent peptides are translocated to the ER post-translationally or co-translationally. This route is dictated by the secretion signal located at the N-terminus of the secreted protein. Within the ER lumen, proteins are modified through folding and gluco/mannosylation. Mis- and unfolded proteins are subject to ERAD, in which they are retro-translocated from the ER to the cytoplasm and degraded by the proteasome. (3) Protein trafficking from the ER to the Golgi and modification within the Golgi. Properly folded/modified proteins are transported to the Golgi via COPII vesicles. Retrograde transport within the Golgi and from the Golgi to the ER is regulated by COPI vesicles. Proteins are further gluco/mannosylated in the Golgi. (4) Protein trafficking between the Golgi and vacuole and exocytosis. Secretory proteins are trafficked from the Golgi to the exterior through vesicles recognized by receptors found on the cell membrane. Secreted proteins are often degraded by proteases secreted into the culture medium. The large retromeric complex regulates protein missorting from the Golgi to the endosome, which later develops into the vacuole. Proteins sorted to the vacuole are subject to degradation by vacuolar proteases.

In this study, we adopted a multi-factorial approach to improve heterologous protein secretion in *Sb*. We first constructed an *Sb* strain that secretes a *C. difficile* Toxin A neutralizing peptide (NPA) and measured NPA secretion kinetics over time. We next investigated the effect of copy number on NPA secretion through the use of different plasmidic and genomically integrated expression cassettes. We also investigated how the mode of ER translocation affects the NPA secretion by testing several native and synthetic secretion signals. We finally engineered *Sb*’s secretory pathway itself, exploring (i) expansion of ER by modifying lipid composition, (ii) disruption of QC machinery in the ER to eliminate ERAD, (iii) minimisation of intra-organelle trafficking and mannosylation modifications in Golgi, (iv) optimization of post-golgi trafficking to eliminate vacuolar sorting and (v) elimination of proteases to diminish vacuolar and extracellular degradation. These modifications were enabled through use of genome editing and proteomics-guided combinatorial strain engineering. Overall, these efforts generated a library of engineered *Sb* strains with a variety of single and combinatorial knockouts to the secretory pathway, collectively leading to *Sb* strains with >10-fold higher NPA secretion than wild-type.

## 2. Methods and Materials

### 2.1. Strains and Culture Media

*Escherichia coli* NEB Stable, NEB 5α, and NEB 10β were used for plasmid construction and maintenance. *E. coli* cells were grown in lysogeny broth (LB) (5g/L yeast extract, 10 g/L tryptone, 10g/L NaCl) at 37 °C supplemented with ampicillin (100 μg/mL), kanamycin (50 μg/mL) or chloramphenicol (34 μg/mL), as appropriate. *Saccharomyces boulardii* ATCC-MYA796Δ*ura3* was used to construct NPA-secreting strains (Jensen et al., 2022). Yeast cultures for plasmid transformation and genome editing were grown in yeast extract-peptone-dextrose (YPD) medium (50 g/L YPD Broth (Sigma-Aldrich)). For biomass generation prior to secretion experiments, yeast cultures were grown in synthetic complete media (CSM) (pH 4.25) containing 0.67% (w/v) Yeast Nitrogen Base Without Amino Acids (Sunrise Science Products), 0.77 g/L Yeast Synthetic Media Dropout Mix (uracil, i.e., CSM-U+1XAA, Sunrise Science Products), and glucose (2% (w/v), Sigma Aldrich). For peptide secretion experiments, yeast cultures were grown in synthetic complete media (pH 7.04) containing 0.67% (w/v) Yeast Nitrogen Base Without Amino Acids (Sunrise Science Products), 1.54 g/L Yeast Synthetic Media Dropout Mix (uracil, i.e. CSM-U+2XAA, Sunrise Science Products), 20.4 g/L potassium phthalate salt (Sigma Aldrich), and glucose (2% (w/v)) (Sigma Aldrich).

### 2.2. Plasmid and Strain Construction (Table S4)

All strains (Supplementary Table S1), oligos (Supplementary Table S2) and gene fragments or gBlocks (Supplementary Table S3) used in this study are in the supplementary information. An entry plasmid (DD183) for peptide secretion was constructed using Gibson cloning (Gibson et al., 2009). The backbone was amplified from another plasmid (ISA002) consisting of a yeast promoter (*pTDH3*), a yeast terminator (*tTDH1*), a yeast selective marker (URA3), a yeast origin of replication (2*μ*) and an *E. coli* selective marker and origin of replication. The GFP insert for this entry plasmid was amplified from DDgb024 (a gene fragment synthesized by IDT) consisting of α mating factor secretion signal sequence, 6xHis-tag sequence, *E. coli* sfGFP expression cassette (BBa_J72163_GlpT_promoter, sfGFP_RBS, and sfGFP - BBa_B0015_terminator) and myc tag sequence. The NPA sequence was inserted in the entry plasmid by replacing the *E. coli* sfGFP expression cassette via Q5 site-directed mutagenesis (New England Biolabs) following the manufacturer’s protocol, leading to the reference secretion plasmid, DD224.

The vectors with other secretion signals were also constructed using Gibson cloning. DD224 was amplified as the backbone (omitting the α mating factor), and the other secretion signal sequences were ordered as gene fragments by IDT and amplified, generating DD226 (SUC1, DDgb026), DD226 (α mating factor, pre, DDgb025), DD280 (Yap3-TA57, DDgb027), DD366 (preOST1-proαMF (I), DDgb028) and DD367 (preOST1-proαMF (MUT1), DDgb029). The NPA secretion vector (DD489) with a low copy origin was constructed using Gibson cloning. DD224 was amplified as the backbone (minus the 2*μ* origin) and the CEN6/ARS4 origin was amplified from the yeast part plasmid (YTK076) from MoClo-YTK (Lee et al., 2015). For all Gibson assembly reactions, each fragment was amplified with tailed primers sharing 20 bp homology. The Gibson Assembly was done according to the manufacturer’s instructions followed by transformation to *E. coli*.

The secretory gene knockouts were constructed via CRISPR-Cas9 genome editing. For each knockout, 1-4 guide RNA (gRNA) - Cas9 plasmids were constructed via Golden gate assembly. gRNA sequences (Table S2) were ordered as complementary single stranded DNA oligos. These were phosphorylated (2 μL 100 μM oligo stock, 2 μL 10X T4 DNA ligase buffer (New England Biolabs), 1 μL T4 PNK (New England Biolabs), 5 μL sterile water, incubated at 37 °C for 1 hour followed by inactivation at 65 °C for 20 minutes), annealed (5 μL phosphorylated forward oligo, 5 μL phosphorylated reverse oligo, 90 μL sterile water) and cloned into a Cas9-gRNA GFP-dropout vector (DD110) via Golden Gate cloning. The Golden Gate reaction mixture contained 0.5 μL of 40 nM of each DNA part (20 fmol), 0.5 μL T7 ligase (EB), 1.0 μL T4 Ligase Buffer (NEB), and 0.5 μL BsaI (10,000 U/mL, NEB), with water to bring the final volume to 10 μL. The NPA secretion cassette for genome integration (DD412) was constructed as a repair template via Golden Gate cloning. Golden Gate assembly protocol was performed on a thermocycler with the following program: 30 cycles of digestion (42 °C for 2 min) and ligation (16 °C for 5 min), followed by a final digestion (60 °C for 10 min) and heat inactivation (80 °C for 10 min). Plasmid ISA1145 provided the gRNA and Cas9 nuclease to integrate the NPA cassette into INT1 (Durmusoglu, Al’Abri, et al., 2021). Repair templates for each gene knockout were ordered as gene fragments from IDT. They consisted of TAATTA basepairs flanked by approximately 250 bp upstream and 250 bp downstream of the target gene.

### 2.3. Yeast Transformations

We used the *Sb* competent cell preparation and transformation protocol from (Durmusoglu, Al’Abri, et al., 2021), which is based on the protocol from (Gietz & Woods, 2002). After transformation, the cells were plated on appropriate growth/selection media. *Sb* strains cultured overnight in appropriate media (Yeast Extract–Peptone–Dextrose (YPD) or Yeast Synthetic Drop-out) in a shaking incubator (37°C, 250 rpm). Saturated cultures were subinoculated to prewarmed media at starting OD 0.25 and grown for 3-4 hours to OD600 1.0 (37°C, 250 rpm). Cells were then harvested at 3000g for 5 min at room temperature and washed twice with sterile water and third time before with 100 mM Lithium Acetate. Supernatant was then removed. Then, solutions were added to the cells in the following order: 260 μL 50% PEG3350 (Fisher Scientific), 36 μL 1 M Lithium Acetate (Sigma-Aldrich), 50 μL of 2 mg/mL single-stranded salmon sperm DNA (Invitrogen, 15632011), 0.1–10 μg DNA. The mixture was then gently vortexed and then incubated at 42 °C for 1 hr. This mixture was then centrifuged at 3000g for 1 min and the supernatant was discarded. The cell pellet was resuspended in 1 mL YPD by gently pipetting up and down and this tube was placed at a shaking incubator (37°C, 250 rpm) for 1 hr. Then, the cell suspension was centrifuged for 1 min at 3000g, resuspended in 25 μL sterile water, and plated on an appropriate growth media.

### 2.4. High-throughput Strain Screening

Once the strains were constructed via genome editing and/or transformation, 3-5 colonies (i.e., clones) were picked for each strain and re-struck on CSM-URA plates, to obtain enough biomass to start stock cultures for high-throughput cultivation experiments. These plates were incubated at 37°C for 2 days. 3 colonies from each streak were inoculated in 1 mL CSM-URA (1XAA, pH 4.25) and grown overnight at 37°C, 250 rpm for 16-18 hours. Then, each culture was washed twice with sterile water and resuspended in 1 mL CSM-URA (2XAA, pH 7.04, supplemented with potassium phthalate (20.4 g/L)) and further diluted in the same media to a final cell density of OD 3. 50 μL of the cell suspension was used as seed culture for the high-throughput cultivations.

High-throughput cultivations were conducted in Biolector II, High-Throughput Microfermentation Platform (m2p Labs). For each strain, each clone (i.e., colony from the original transformation) was cultured in 3 replicate microfermentations. Microfermentations were conducted in a 48-well flower plate (MTP-48-B, without optodes + biomass, fluorescence) in 1.5 ml CSM-URA (2XAA, pH 7.04, supplemented with potassium phthalate (20.4 g/L)) at a starting cell density of OD 0.1. Flower Plates were incubated in the Biolector II at 37°C and 85% humidity with 1000 rpm shaking for 12-48 hours for the time course experiment involving the NPA reference strain, 36 hours for secretion signal experiments, 48 hours for copy number experiments, 48 hours for the gene knockout experiments and 48 hours for the combinatorial experiments.

### 2.5. Detection of NPA Secretion via SDS-PAGE

Supernatants from the NPA secretion reference strain (Sb-NPA) were acetone precipitated by mixing 1 volume of the supernatant with 4 volumes of pure acetone. The mixtures were incubated overnight at −20 °C. Protein pellets were collected by centrifugation at 5,000g for 5 minutes. Protein pellets were then resuspended in 200 μL 1X Tricine Sample Buffer (Bio-Rad) and 20 μL of protein sample was loaded to 4-20% MiniProtean Tris-Tricine Precast gels (Bio-Rad). 10 μL Precision Plus Dual Xtra Protein Standard was also loaded as a ladder. The gels were run for 30 minutes at 200V. The gels were fixed in a 40% methanol, 10% acetic acid and 50% water mixture for 30 minutes, stained with Imperial protein stain (Thermo Scientific) for 4 hours, and washed in deionized water overnight on a rocking mixer. The gels were imaged using a BioRad GelDoc EZ system.

### 2.6. Isolation of NPA via His-Tag Isolation and Pulldown

Acetone-precipitated Sb-NPA supernatants were resuspended in 1 mL 1X binding/wash buffer (50 mM sodium phosphate, 300 mM NaCl, 0.01% Tween-20). This solution was incubated with 50 μL Dynabeads (Invitrogen) in a microcentrifuge tube on a rotating roller for 10 minutes. The beads were collected using a magnet and were washed 4 times with 300 μL binding/wash buffer gently. The beads were collected using a magnet and the supernatant was discarded. The beads were resuspended in 100 μL elution buffer (300 mM imidazole, 50 mM sodium phosphate, 300 mM NaCl, 0.01% Tween-20) and incubated on a rotating roller for 10 minutes. The beads were removed by a magnet and the eluate was analyzed via SDS-PAGE.

### 2.7. Detection of NPA Secretion via ELISA

Supernatants from microfermentations were diluted in 1xTBS with 0.5% Tween-20 buffer, Similarly, a multi-tag protein standard (Genscript) was serially diluted in 1xTBS with 0.5% Tween-20 buffer, with concentrations ranging between 4 mg/L - 0.0625 mg/L. 100 μL protein standard and supernatant dilutions were incubated in Pierce Nickel Coated Plates (Thermo Scientific) for 1 hour on an orbital shaker at 125 rpm. The plates were then washed 3 times with 200 *μ*L 1xTBS with 0.5% Tween-20. A primary antibody (c-myc tag Polyclonal Antibody, HRP, Bethyl Laboratories) solution was prepared at 1:10000 dilution in 1xTBS with 0.5% Tween-20 buffer. 100 *μ*L of this diluted antibody solution was added to the wells and the plates were incubated for 45 minutes on an orbital shaker at 125 rpm. The plates were then washed 3 times with 200 *μ*L 1xTBS with 0.5% Tween-20. 100 *μ*L 1-Step^™^ Ultra TMB-ELISA Substrate Solution (3,3’,5,5’-tetramethylbenzidine) (ThermoFisher) was added to the plates and incubated for 1 minute to start HRP activity and 100 *μ*L 2M sulfuric acid was added to stop the reaction. Absorbance of each well was detected at OD450 using a SpectraMax iD3 Multi-Mode Microplate Reader (Molecular Devices). Among the 3-5 colonies screened for each strain, the clone with best growth and highest NPA secretion titers was chosen for data analysis on GraphPad. During the data analysis protein standard concentrations and NPA were normalized using their respective molecular weights.

### 2.8. Sample Preparation for Proteomics

Sb-NPA was cultured in 5 mL CSM-URA (1XAA, pH 4.25) at 37°C for 72 hours. Supernatants from the cultures were acetone precipitated by mixing 1 volume of the supernatant with 4 volumes of pure acetone. The mixtures were incubated overnight at −20 °C. Protein pellets were collected by centrifugation at 5,000g for 5 minutes and further processed for proteomics.

### 2.9. Modified Filter Aided Sample Preparation (FASP) for Proteomics

Samples were thawed, lyophilized, and reconstituted in 100 μL 50 mM ammonium bicarbonate (NH4HCO3 or ABC). The content of protein in the samples was determined with the A280 Total Protein Assay (Thermo Scientific, Wilmington, DE). A volume equivalent to 100 μg protein per sample was taken through a modified filter-aided sample preparation (FASP) protocol with tryptic digestion for bottom-up proteomics. Briefly, the protein from each sample was diluted with 50 mM ABC to arrive at a final volume of 200 μL (0.5 μg/μL). For reduction of protein disulfide bonds, 20 mM dithiothreitol (DTT) in 50 mM ABC was added to each sample to arrive at a final DTT concentration of ~5 mM. The samples were incubated for 30 min at 60°C and then allowed to cool to room temperature (~ 5 min) before proceeding to the next step. A 500 mM iodoacetamide (IAA) solution (in 50 mM ABC) was added to each sample to arrive at a final IAA concentration of ~15 mM. The reaction was allowed to proceed in the dark at room temperature for 20 min. Each sample solution was then transferred onto separate, passivated (with 20 μL of 50 mM ABC in 1% sodium deoxycholate or SDC) Vivacon 500 3 kDa molecular weight cutoff (MWCO) filtration units from Sartorius Stedim Biotech, Goettingen, Germany (of note, 2 kDa MWCO filtration units were also tested, yielding similar results). The samples were then centrifuged at 12500g for 25 min and the flowthrough was discarded. Reduction and alkylation of the sample protein was followed by a series of washing steps. A 200 μL volume of 50 mM ABC in 8 M urea was added to each filter and then the filtration units were centrifuged at 12500g for 45 min. This step was repeated one more time to ensure a thorough wash. The flowthrough was discarded from the collection tubes following these washing steps. A 200 μL volume of 50 mM ABC was then added to each filter and the filtration units were centrifuged at 13000g for 45 min. This washing step was also repeated one more time and the flow-through solutions were discarded. After replacing the collection tubes with fresh ones, 80 μL of Trypsin Gold (Promega, Madison, WI) solution (in 50 mM ABC) was added to each filter for an enzyme/substrate ratio of 1:50. The tops of the filtration units were wrapped with parafilm to minimize evaporation and the units were then incubated in an Eppendorf ThermoMixer (Hamburg, Germany) at 37°C and 600 rpm for 6 hrs. Next, the filtration units were then centrifuged at 12500g for 35 min. 15.8 μL of 50 mM ABC was then added to each MWCO filter, followed by centrifugation at 12500g for 20 min, so as to elute any remaining tryptic peptides from the MWCO filters. The filters were discarded and 4.2 μL of 6M hydrochloric acid (HCl) was added to each sample to quench the digestion. The digested samples (100 μL each) were stored at −20°C until analysis by nanoLC-MS/MS.

### 2.10. LC-MS/MS analysis

The protein digests were reconstituted with 100 μL Mobile Phase A (MPA, 2% acetonitrile in water with 0.1% formic acid) and 2 μL injections were analyzed by reversed phase nano-liquid chromatography - mass spectrometry (nano-LC-MS/MS) using an Orbitrap Exploris 480 Mass Spectrometer (Thermo Scientific, Bremen, Germany) interfaced with an EASY nanoLC-1200 system (Thermo Scientific, San Jose, CA, USA). An EASY-Spray nano-flow source (Thermo Scientific, San Jose, CA) effected electrospray ionization of peptide digests. The sample peptides were concentrated, desalted and separated using a ‘trap and elute’ column configuration - consisting of a 0.075 mm × 20 mm C18 trap column with particle size of 3 μm (Thermo Scientific Accclaim PepMap100) in line with a 0.075 mm × 250 mm C18 analytical column with particle size of 2 μm (Thermo Scientific EASY-Spray^™^) - with nanoLC flowrate maintained at 300 nL/min. Peptides were eluted using a 140 min gradient, ramping from 5 to 25% Mobile Phase B (MPB, 80% acetonitrile with 0.1% formic acid) over 105 min, followed by another ramp to 40% MPB over 15 min, and then a steep ramp to 95% MPB in 1 min, at which point MPB was maintained at 95% for 17 min for column washing. Eluting tryptic peptides were ionized by subjecting them to 1.8 kV in the ion source. The ion transfer tube temperature was maintained at 275 ^o^C. The peptides were interrogated by full MS scan and data-dependent acquisition (DDA) MS/MS. Full MS data was collected with an m/z scan range of 375 to 1,600 in positive ion mode at 120 K resolving power with 300% normalized Automatic Gain Control (AGC) Target, 100 ms maximum injection time and RF lens of 40%. MS/MS scans were collected at 7.5 K mass resolving power, with a 1.5 m/z isolation window, 30% normalized Higher-Energy Collisional Dissociation (HCD), 100% normalized AGC Target, custom maximum injection time and dynamic exclusion applied for 20 s periods.

### 2.11. Proteomics Data Interrogation

The raw nanoLC-MS/MS files were interrogated with Proteome Discoverer 2.4.0.305 (PD, Thermo Scientific, San Jose, CA) software against the *Saccharomyces boulardii* protein database on UNIPROT (Khatri et al., 2017). This database were searched with the following parameters: trypsin (full) as the digesting enzyme, a maximum of 2 missed trypsin cleavage sites allowed, 5 ppm precursor mass tolerance, 0.02 Da fragment mass tolerance, dynamic modifications on [a] methionine (oxidation) [b] protein N-terminus (acetyl) [c] protein N-terminus (Met-loss) [d] protein N-terminus (Met-loss+Acetyl), as well as static carbamidomethyl modifications on cysteine residues. The SEQUEST HT algorithm was employed in data interrogation. Percolator peptide validation was based on the q-value, and minimal false discovery rate (FDR) < 0.01 was considered as a condition for successful peptide assignments.

## 3. Results and Discussion

### 3.1. *Sb* can secrete a recombinant anti-toxin peptide

As an exemplary therapeutic, we chose to secrete a peptide (NPA, DYWFQRHGHR) that inhibits the glucosyltransferase activity of *C. difficile* Toxin A (Xiao et al., 2022). Due to their small size, peptides exhibit higher diffusivities than larger proteins (e.g., nanobodies and antibodies), and so are a promising class of therapeutics for *in situ* biomanufacturing. The codon-optimized NPA sequence was tagged at the N-terminus with a 6x-His tag, and at the C-terminus with a myc tag to facilitate peptide detection via ELISA. Direction of tagged NPA to the secretory pathway was initially enabled by the α-mating factor secretion signal (αMF-prepro). αMF-prepro is 89 amino acids (aa) long and contains three domains: a 19 aa pre-region guiding translocation to the ER, a 64 aa pro-region guiding transport from the ER to Golgi, and a 6 aa spacer region containing the KEX2 protease recognition site that is processed in the ER. αMF-prepro has been used for secretory production of a wide range of recombinant proteins in *Sc* (Besada-Lombana & Da Silva, 2019; Z. Liu et al., 2012; Zsebo et al., 1986). The expression of NPA was controlled by the constitutive TDH3 promoter (*pTDH3*) and TDH1 terminator (*tTDH1*) (**Figure S1**), both of which enable high levels of gene expression in *Sb* (Durmusoglu, Al’Abri, et al., 2021; Lee et al., 2015). This expression cassette was placed on a plasmid containing a *URA3* selective marker and high-copy (2*μ*) origin of replication. We chose the episomal (2*μ*) origin and URA3 selective marker, as this combination achieved the strongest plasmidic gene expression in *Sb* (Durmusoglu, Al’Abri, et al., 2021). While genomic integration is often more appropriate for *in situ* biomanufacturing, plasmids allow for rapid prototyping of engineered strains (Hohnholz et al., 2017; Lee et al., 2015). The *Sb* strain containing this NPA expression vector was referred to as Sb-NPA. As a negative control, we constructed another plasmid containing a non-coding “spacer” sequence in place of the “promoter - gene - terminator” cassette on the same expression vector. This strain is referred to as Sb-Spacer.

**Figure S1.**
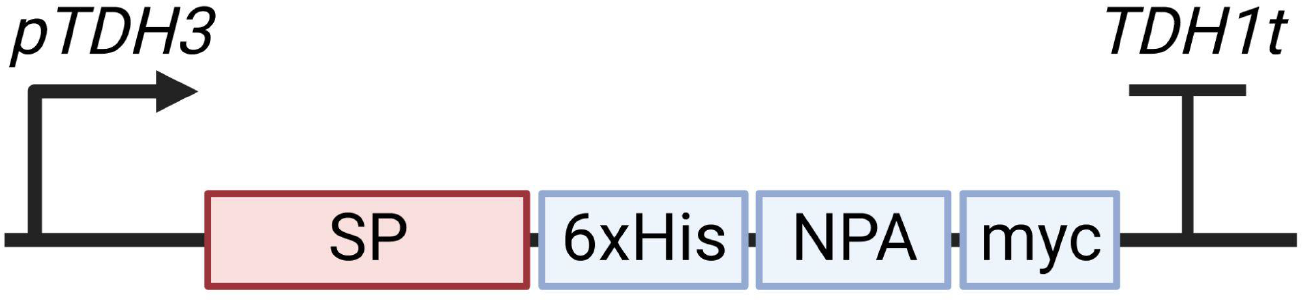
Schematic overview of the expression cassette for secretory production of therapeutic peptide NPA. The expression was regulated by the strong constitutive TDH3 promoter and TDH1 terminator. Native and synthetic signal peptide sequences were cloned upstream of the NPA sequence, which was flanked by a poly-histidine tag and myc tag on its N- and C-terminus, respectively. This cassette was inserted into high- or low-copy plasmids or inserted into the genome.

Sb-NPA and Sb-Spacer were first cultured in triplicate in 5 mL complete synthetic media lacking uracil (CSM-U) with 1X concentration of amino acids (1XAA) at 37 °C for 24 hours. Then, 2 mL culture supernatant was collected, acetone precipitated, and resuspended in tricine buffer. These concentrated supernatants were then run on 4-20% Tris-Tricine SDS-PAGE gels. Based on coomassie blue staining, Sb-NPA and Sb-Spacer did not secrete the peptide to the culture medium (**Figure S2a**). We hypothesized that low secretion levels could be due to the media formulation, as previous studies on secretory production of proteins in *Sc* utilized complete synthetic media containing 2XAA (Hou et al., 2012b; Huang et al., 2018). When we cultured the strains in CSM-U+2XAA at 37 °C for up to 72 hours, we observed a large increase to the levels of secreted proteins (**Figure 2**, **Figure S2b-c**). We first observed several high molecular weight bands that were present in both supernatants, which could comprise proteases, phosphatases and/or glycosyl hydrolases, all of which are known components of the *Sc* secretome (Smeekens et al., 2017). We also observed a band in Sb-NPA lanes that was not present in the Sb-Spacer lanes (**Figure 2a**). This band was 2-5 kDa in size, corresponding to the predicted molecular weight of tagged NPA (3.6 kDa). In order to confirm whether this additional band was indeed tagged NPA, we purified acetone-precipitated supernatants using nickel beads, which selectively bind proteins with polyhistidine tags. We ran His-tag isolated samples from Sb-NPA cultured for 24 hours and 48 hours on SDS-PAGE gels and observed bands with sizes consistent with tagged NPA (**Figure S2d-e**), further supporting the presence of NPA in the supernatant.

**Figure 2.**
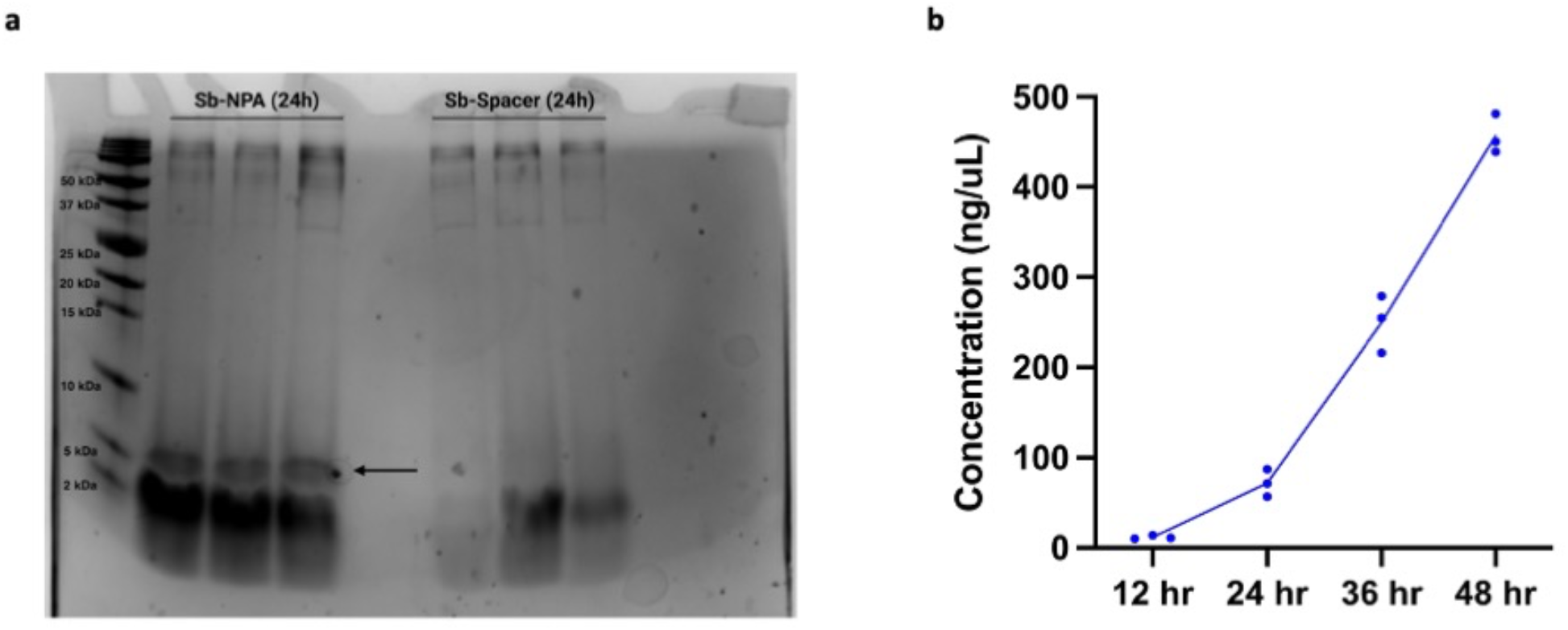
NPA secretion in *Sb*. **(a)** SDS-PAGE image of precipitated supernatants of NPA secretor (Sb-NPA) and non-expressing (Sb-Spacer) strains. Each strain was cultured in triplicates in culture tubes at 37 °C for 24 hours. The black arrow indicates the band corresponding to tagged NPA. **(b)** Secreted NPA titers by Sb-NPA over 48h in a BioLector microfermenter. Dots represent the NPA concentration for each replicate cultivation and lines connect the average NPA concentration for each timepoint.

**Figure S2.**
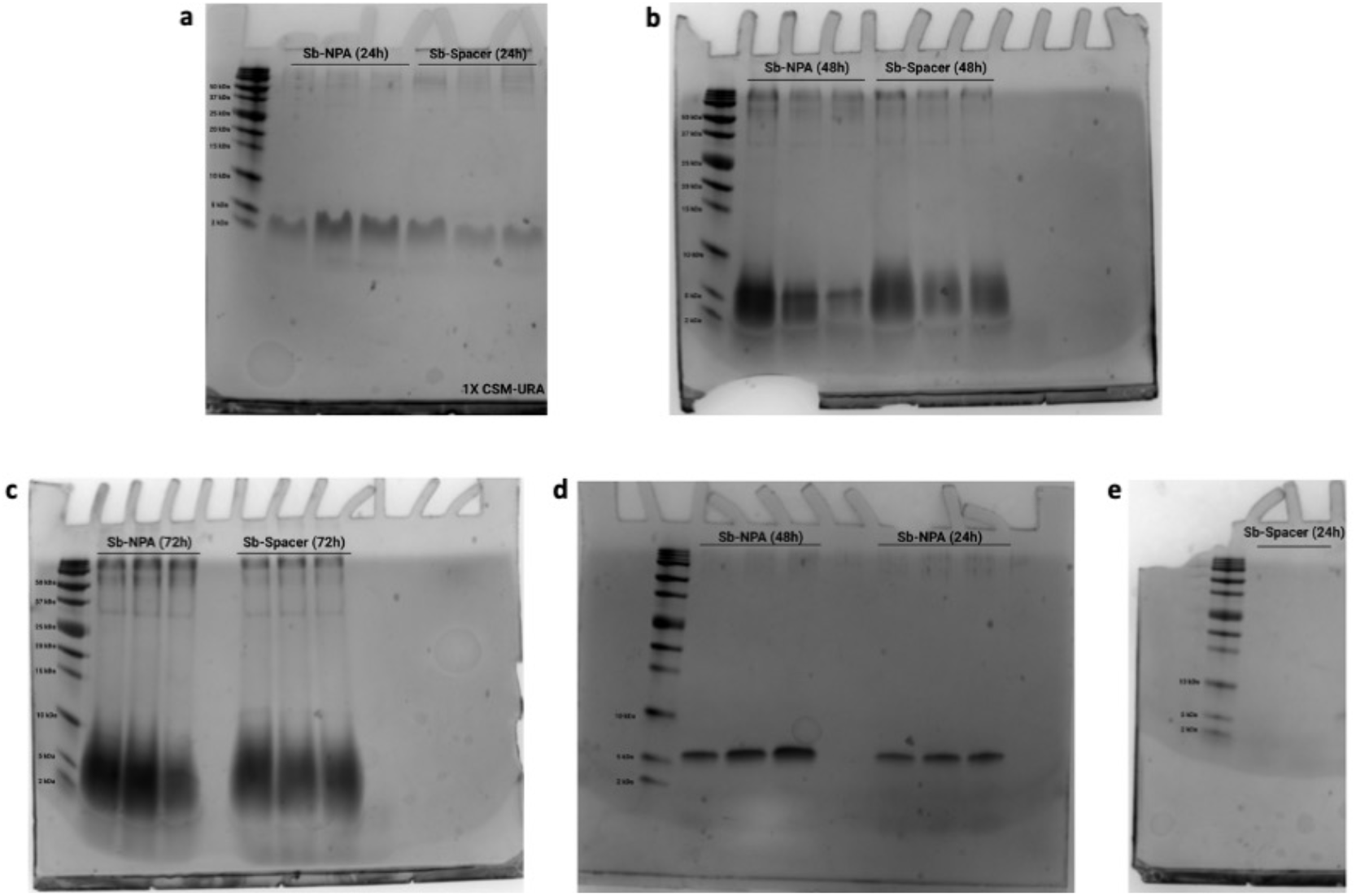
Detection of NPA secreted in culture media via SDS-PAGE. (a) Sb-NPA was cultured in (1X) CSM-URA for 24 hours and supernatants were precipitated and run on 4-20% Tris-Tricine gels. Sb-NPA and Sb-Spacer supernatants were collected after 48 hours (b) and 72 hours (c), precipitated, and run on 4-20% Tris-Tricine gels. Each strain was cultured in triplicate in culture tubes at 37 °C. Precipitated Sb-NPA (d) and Sb-Spacer (e) supernatants from 24 hours (right 3 lanes) and 48 hours (left 3 lanes) were processed through Dynabeads His-Tag Isolation and Pulldown to isolate poly-histidine tagged NPA present in the precipitates. Eluates were run on 4-20% Tris-Tricine gels.

Next, we sought to understand how cultivation duration impacts NPA secretion. On SDS-PAGE images from 48-hour cultures, NPA bands were visible but partly obscured by other low molecular weight products (**Figure S2b**). This phenomenon was further pronounced in samples from 72-hour cultivations, preventing visual assessment of NPA concentration using SDS-PAGE (**Figure S2c**). It is known that native proteases can degrade secreted recombinant proteins over long-term culture, potentially leading to the peptide-sized fragments we observed (Landowski et al., 2015). In order to quantify NPA concentrations more accurately, we utilized an enzyme-linked immunosorbent assay (ELISA) specific to both 6x-His and myc tags. Specifically, NPA was bound to a nickel-coated plate via its 6x-His tag and quantified via labeling with an anti-myc antibody conjugated to horseradish peroxidase (HRP). As such, only intact NPA is detected via this assay. Sb-NPA was cultured in triplicate in 1.5 mL CSM-U+2XAA at 37 °C for 12, 24, 36 and 48 hours in a Biolector II microfermentor. 20 μL supernatants were analyzed via ELISA (Figure 2b). At 12 hours, NPA concentration in supernatant was 14 mg/L, and at 24 hours its concentration was 71 mg/L. The rate of NPA accumulation in the supernatant increased after 24 hours, leading to 255 mg/L and 452 mg/L of NPA in the supernatant after 36 hours and 48 hours of cultivation, respectively. This increase could be attributed to increased secretory activity of the cell as it approached stationary phase, a known phenomenon in other yeast strains (Huang et al., 2017). Thus, we were able to achieve secretory production of anti-toxin peptide NPA in *Sb*, reaching a maximum of titer 452 mg/L. To our knowledge, this is the highest recombinant protein secretion titer achieved in wild-type *Sb*.

### 3.2. Peptide secretion is correlated with gene copy number

Plasmids are commonly used to encode recombinant proteins due to their ease of manipulation and high copy number. However, genomic integration of synthetic DNA allows for stable gene expression in the absence of artificial selection pressures, and is particularly relevant for *in situ* biomanufacturing. We therefore asked whether NPA copy number would affect secretion, hypothesizing that higher copy numbers would result in higher secretion levels. We thus constructed 2 additional NPA secretion vectors: (i) a plasmid vector with the centromeric (CEN6/ARS4) origin and (ii) a genomic integration vector targeted to the noncoding integration locus (INT1) enabling a high expression level in *Sb* (*Durmusoglu, Al’Abri, et al., 2021*). We cultured these two new NPA-secreting *Sb* strains in triplicate in 1.5 mL CSM-U+2XAA at 37 °C for up to 48 hours in a Biolector II microfermentor (**Figure 3**). *Sb* strains with genomic integration achieved an NPA titer of only 76 mg/L. Using a plasmid with a centromeric origin improved NPA concentration 2.95-fold, reaching 224 mg/L in the supernatant. As before, utilizing a plasmid with an episomal origin enabled high NPA secretion levels (458 mg/L), 6-fold and 2-fold higher than encoding NPA on the genome or a centromeric vector, respectively. Plasmids with the URA3 selective marker with centromeric origins are maintained at 4-8 copies per cell, while those with episomal (2μ) origins are maintained at 28-58 copies per cell (Karim et al., 2013). Given that we integrated NPA into both copies of the *Sb* genome, genomic integration yielded 2 copies per cell. These differences in copy number are thought to modify mRNA levels, in turn impacting translation and secretion rates (Besada-Lombana & Da Silva, 2019; Z. Liu et al., 2012).

**Figure 3.**
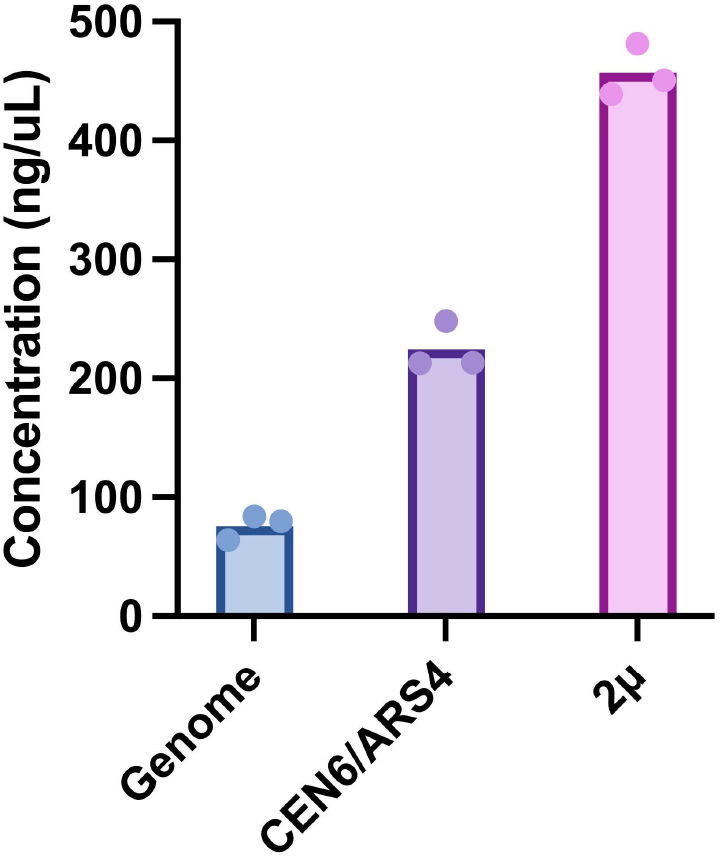
Effect of gene copy number on peptide secretion in *Sb*. NPA with αMF secretion signal was placed on the genome or on plasmids with low-copy (*CEN6/ARS4*) or high-copy (2μ) yeast origins, and NPA secretion was measured after 48h. Bars represent the average NPA concentration across three cultivations, and dots represent the NPA concentration in each cultivation for a given strain.

Gene copy number can also impact cell fitness. Although multicopy plasmids can often cause metabolic burden in yeast, resulting in growth deficiency (Rouches et al., 2022), their impact on cell fitness was rather small compared to the significant improvement in secreted NPA amount (**Figure S3**). Therefore, it will be important to explore the relationships between NPA copy number, *Sb* colonization levels, and *in vivo* peptide titer in future work.

**Figure S3.**
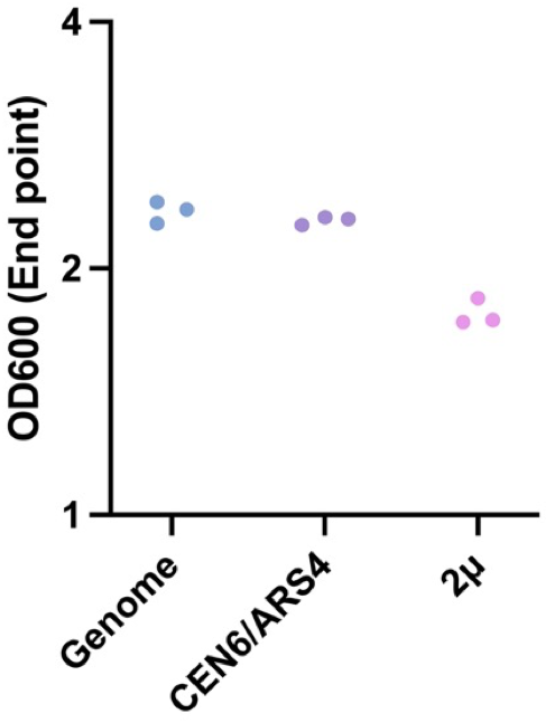
Final cell density of NPA secretor strains with varying copy numbers. Each strain was cultured in triplicate in a Biolector II microfermentor for 48 hours at 37 °C. OD600 values were obtained at the end of the cultivation via spectrophotometer.

### 3.3. Hybrid secretion signal sequences improve peptide secretion

The first step in the secretory pathway is transfer of the peptide chain into the ER lumen through the ER membrane. This transfer, i.e., translocation, is directed by the secretion signal fused to the protein destined to be secreted. Secretion signals direct translocation through one of two mechanisms: (i) post-translational translocation or (ii) co-translational translocation. In the former, the mature peptide chain is inserted into the ER lumen after translation is finished. In the latter, the nascent peptide chain is recognized by signal recognition particles (SRPs) found in the cytosol. Once recognized, translation stalls and the ribosome-peptide-SRP complex is translocated to the ER membrane, pushing the nascent peptide to the ER lumen. Even though both processes initiate secretion, cotranslational translocation is thought to further promote the availability of the nascent protein for secretory processes (Jan et al., 2014). The rationale for this is that the secondary and tertiary structures of mature proteins established in the cytoplasm may hinder their translocation across the ER membrane in posttranslational translocation. Also, the ability of each signal to direct secretion changes depending on the protein (e.g., its size, structure, and solubility) it is fused to. Therefore, we wanted to understand the effect of secretion signals on NPA secretion.

We constructed *Sb* strains secreting NPA via 3 native and 3 synthetic secretion signals (**Figure 4**). In the reference strain (Sb-NPA), we utilized *Sb’s* endogenous αMF-prepro secretion signal. This signal has been extensively used in other yeasts to secrete a range of proteins (Aggarwal & Mishra, 2020; Brake et al., 1984; Møller et al., 2017; Wang et al., 2021). Using the αMF-prepro signal, we achieved 250 mg/L NPA after 36 hours of cultivation. Previous work showed that using only the pre-region of αMF is sufficient to achieve secretion in *Pichia pastoris (Pp*) (Fitzgerald & Glick, 2014). When we cultivated *Sb* expressing NPA with only the αMF-pre signal, the concentration of NPA in supernatant was reduced to 132 mg/L. Similar differences between αMF-prepro and αMF-pre were also observed in *Sc* expressing human insulin-like growth factor and *Pp* expressing msGFP. In these works, including the pro-region was found to improve trafficking and processing of recombinant protein through the ER and Golgi (Chaudhuri et al., 1992; Fitzgerald & Glick, 2014). However, the reduction in secretion by the αMF-pre-only strain improved final cell density, presumably due to reduced metabolic burden (**Figure S4**).

**Figure 4.**
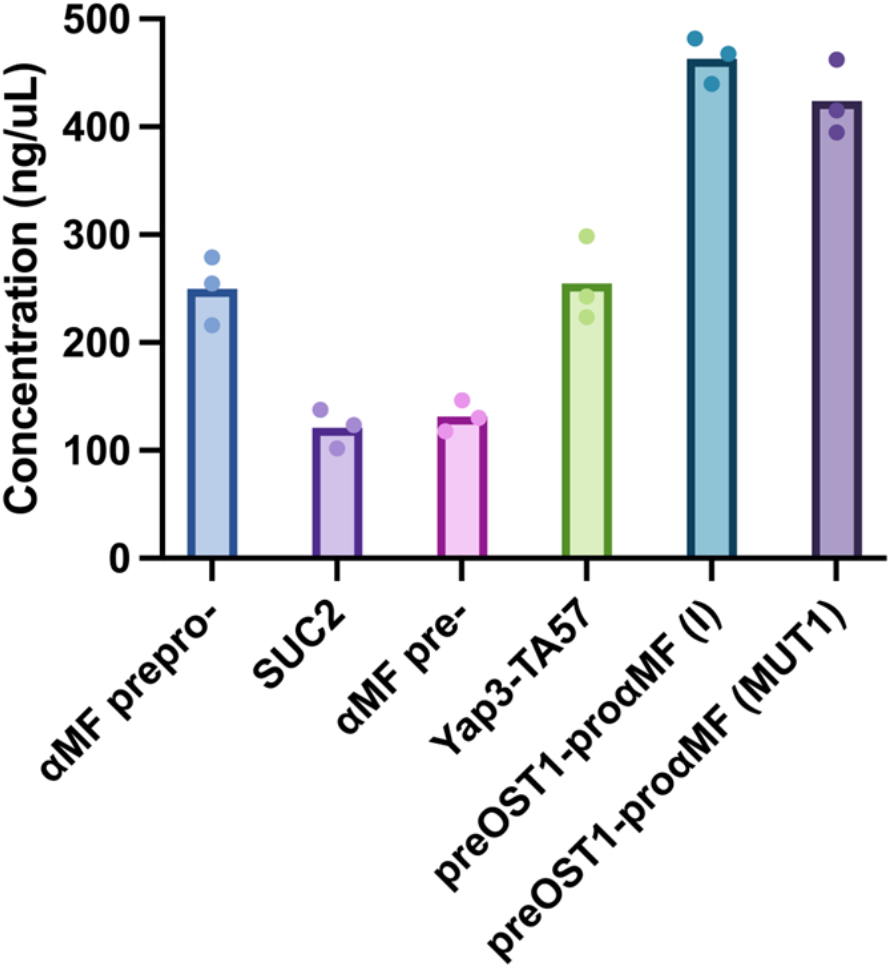
Effect of secretion signal on peptide secretion in *Sb*. NPA secretion cassettes with 3 native and 3 synthetic secretion signals were cloned into a backbone with a high-copy (2μ) origin. Each strain was cultured in triplicate in FlowerPlates in a Biolector II for 36 hours at 37 °C. Bars represent the average NPA concentration across three cultivations, and dots represent the NPA concentration in each cultivation.

**Figure S4.**
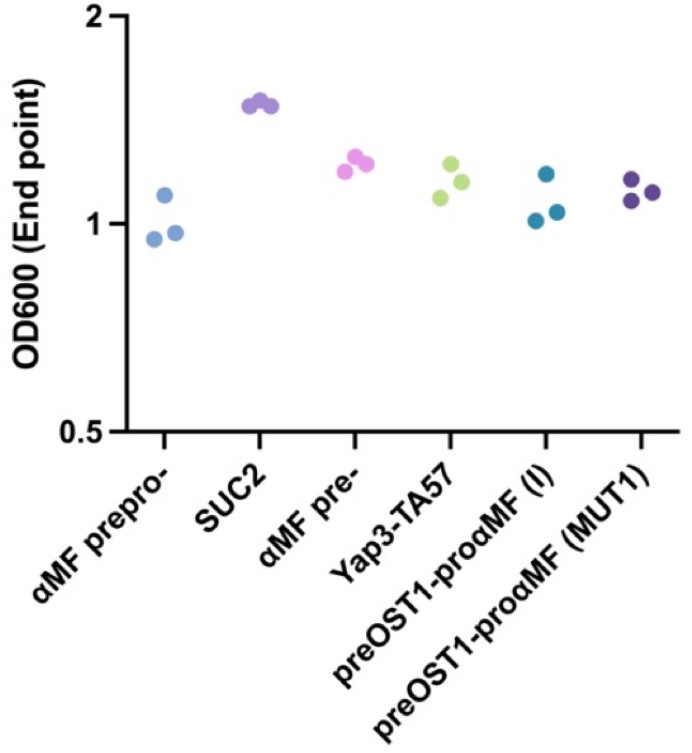
Final cell density of NPA secretor strains with varying secretion signals. NPA secretion cassettes with 3 native and 3 synthetic secretion signals were cloned into a backbone with a high-copy (2μ) origin. Each strain was cultured in triplicate in FlowerPlates in a Biolector II for 36 hours at 37 °C. Bars represent the average endpoint cell density across three cultivations, and dots represent the endpoint cell density in each cultivation.

As the third native secretion signal, we investigated that of invertase (*SUC2*). Invertase is a naturally secreted enzyme that catalyzes conversion of sucrose to fructose and glucose. In *Sc*, the wild-type signal and its mutants have been used to secrete complex recombinant proteins such as human interferon and α-amylase (Chang et al., 1986; Nishizawa et al., 1989; Rothe & Lehle, 1998). In *Sb*, the *SUC2* signal peptide enabled an NPA titer of 121 mg/L, the lowest od the 6 signals we tested. This decreased secretion with the *SUC2* signal could be due to the lack of a pro sequence, thereby impairing the trafficking and processing of NPA in the secretory pathway, similar to the αMF-pre signal.

In addition to native signal peptides, we tested three synthetic signal peptides that were developed either as hybrids between native signals, or as point mutants thereof. These peptides have been reported to improve secretion compared to their native counterparts (Aza et al., 2021; Bose, 2021; Lin-Cereghino et al., 2013). First, we tested a synthetic signal called Yap3-TA57 which was developed to improve the secretion of insulin precursor (IP) (Kjeldsen et al., 1998, 1999b)). This secretion signal consists of a 21 aa pre-region from Yap3p and a 44 aa synthetic pro-region with mutated residues lacking N-linked glycosylation and providing improved enzymatic processing in Golgi. (Kjeldsen et al., 1998, 1999b). Using Yap3-TA57 enabled 255 mg/L NPA secretion, which is similar to what we observed with the αMF-prepro signal. By contrast, in *Sc*, Yap3-TA57 improved the secretory IP yield 81% and secretory α-amylase yield 10% compared to the αMF-prepro signal (Kjeldsen et al., 1998; Z. Liu et al., 2012), showing that signal performance may vary from species to species, similar to what was observed previously between *Sc* and *Pp* for the same synthetic leader (Kjeldsen et al., 1999a).

Since *Sc* and *Pp* can use the same secretion signals to secrete proteins, we wanted to test 2 synthetic secretion signals (preOST1-proαMF (I) and preOST1-proαMF (MUT1)) developed previously in *Pp*. These two sequences contain the pre-region from oligosaccharyl transferase complex (OST1). Unlike the αMF-pre, which enables posttranslational translocation, OST1-pre enables co-translational translocation, potentially improving trafficking efficiency in the early secretory pathway. Both signals contain a mutated αMFpro-region. preOST1-proαMF (I) contains L42S and D83E mutations, whereas preOST1-proαMF (MUT1) contains only L42S (Barrero et al., 2018). Of these, L42S was found to be sufficient for improving protein trafficking by eliminating protein aggregation in the *Pp* ER lumen. When we tested these two signals in *Sb*, preOST1-proαMF (I) achieved slightly more secretory NPA production (463 mg/L) than preOST1-proαMF (MUT1) (424 mg/L). A similar trend was observed in *Pp* for these two variants, confirming that the L42S mutation is more important than D83E for improved secretion. Interestingly, although several synthetic leaders improved secretion compared to Sb-NPA, the cell growth was not substantially impacted (**Figure S4**). In fact, all 3 synthetic signals enabled final cell densities similar to Sb-NPA. Potentially, improved protein trafficking and processing could compensate for the metabolic burden arising from the secretory production of NPA. Overall, by using different secretion signals, we were able to cover a 3.82-fold range of NPA secretion in *Sb*. In the future, more secretion signals, both native and synthetic, with improved secretory profiles could be discovered through genome-wide screening, directed evolution, and machine learning technologies (Aza et al., 2021; Bae et al., 2015; Mori et al., 2015; Z. Wu et al., 2020)(Aza et al., 2021; Bae et al., 2015; Mori et al., 2015; Z. Wu et al., 2020), to expand the toolbox for engineered protein secretion in *Sb*.

### 3.4. Removing vacuolar/extracellular proteases de-bottlenecks peptide secretion

Next, we sought to understand the effect of modifying key steps across the secretory pathway on recombinant peptide yield. We prioritized 4 biomolecular processes involved in the secretory performance (**Table 1**): (i) lipid composition & stress responses, (ii) ER associated degradation (ERAD), (iii) trafficking and modifications within the Golgi, and (iv) vacuolar/extracellular protein degradation. These processes were studied previously in *Sc* (via gene deletion or overexpression), and their effects (improved or deleterious) on recombinant protein secretion have been reported in numerous studies (Huang et al., 2018; Madhavan et al., 2021). We focused on 13 genes regulating the aforementioned processes, and we investigated their effect on secretion performance through CRISPR-Cas9 genome editing, where we generated homozygous deletions in each of these 13 genes in *Sb*. We designed 2-4 guide RNAs (gRNAs) per gene and transformed *Sb* with (i) gRNA-Cas9 plasmid to assist DNA cleavage and (ii) double stranded DNA containing 500 bp of upstream and downstream homology to assist homologous recombination, thereby removing the entire ORF from both chromosomes.

**Table 1.**
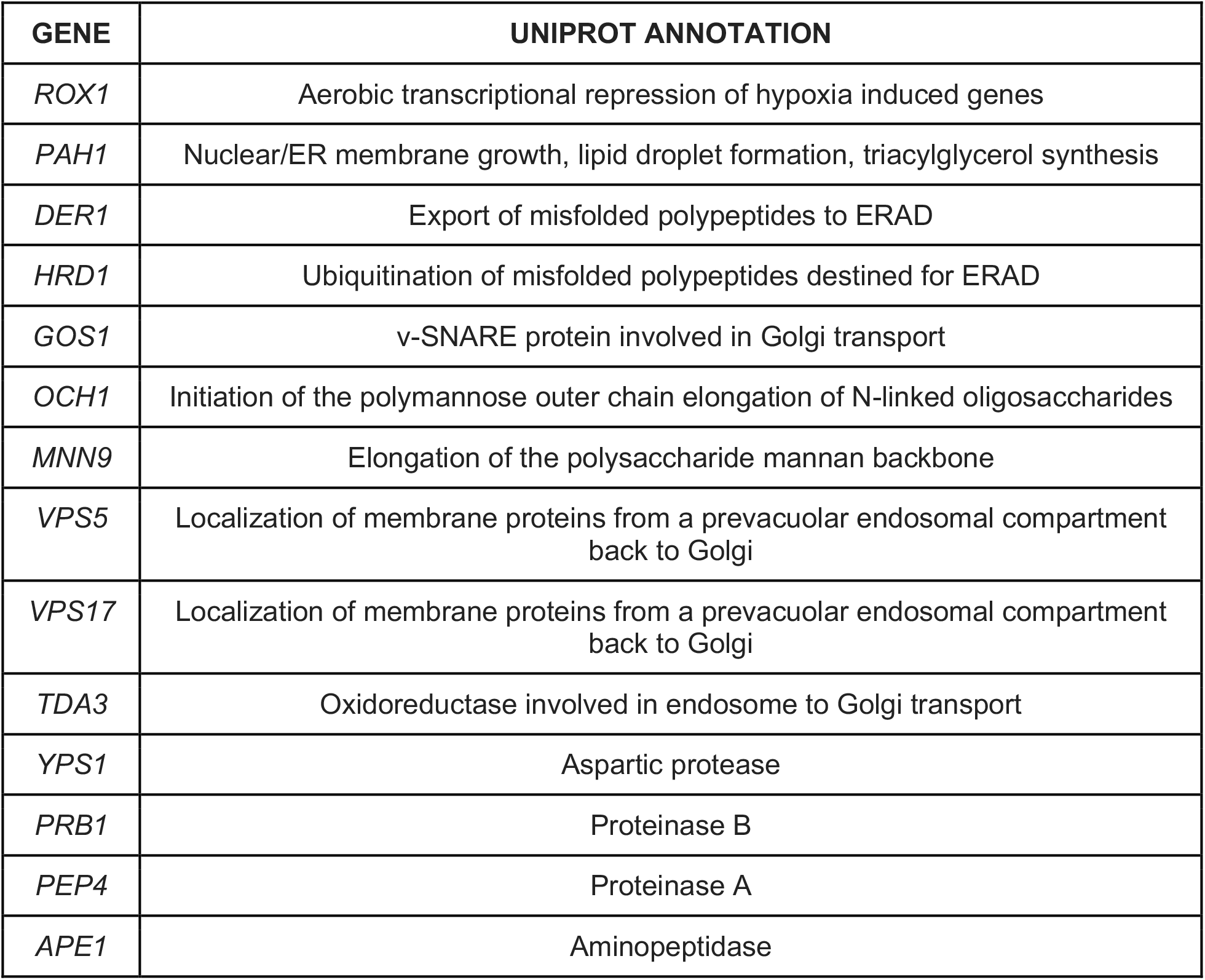
List of genes deleted to optimize secretory pathway in this study *SNARE: Soluble N-ethylmaleimide-Sensitive Factor Attachment Protein Receptor.

| GENE | UNIPROT ANNOTATION |
| --- | --- |
| *ROX1* | Aerobic transcriptional repression of hypoxia induced genes |
| *PAH1* | Nuclear/ER membrane growth, lipid droplet formation, triacylglycerol synthesis |
| *DER1* | Export of misfolded polypeptides to ERAD |
| *HRD1* | Ubiquitination of misfolded polypeptides destined for ERAD |
| *GOS1* | v-SNARE protein involved in Golgi transport |
| *OCH1* | Initiation of the polymannose outer chain elongation of N-linked oligosaccharides |
| *MNN9* | Elongation of the polysaccharide mannan backbone |
| *VPS5* | Localization of membrane proteins from a prevacuolar endosomal compartment back to Golgi |
| *VPS17* | Localization of membrane proteins from a prevacuolar endosomal compartment back to Golgi |
| *TDA3* | Oxidoreductase involved in endosome to Golgi transport |
| *YPS1* | Aspartic protease |
| *PRB1* | Proteinase B |
| *PEP4* | Proteinase A |
| *APE1* | Aminopeptidase |

We first investigated how deleting genes involved in lipid composition and stress response would change NPA secretion. Lipids comprise the organelle and vesicle membranes where important secretory processes take place, and therefore modifying lipid composition could impact secretion efficiency. Rox1p, a transcriptional regulator that controls hypoxia responses under oxidative stress conditions in yeast, was also found to affect lipid composition by affecting lipid and sterol biosynthesis (Jordá & Puig, 2020; L. Liu et al., 2015). Deletion of *ROX1* improved secretory production of hIP and α-amylase in Sc (Jordá & Puig, 2020; L. Liu et al., 2015). This increased production was attributed to changes in lipid composition, particularly to increased ergosterol concentration, which is a significant component of secretory vesicles. *Sb*Δ*rox1* increased secretory NPA production 40% (**Figure 5**). However, this improvement caused a reduced final cell density (**Figure S5**). We speculate that this could be due to a reduced cell size (thereby reducing the culture optical density), because deletion of *ROX1* results in *Sc* cells with smaller size (Liu et al. 2015). PAH1p is a phosphatidate phosphatase found in the cytoplasm and is responsible for diacylglycerol synthesis from phosphatidic acid found in the ER membrane. Deletion of *PAH1* results in impaired lipid droplet formation and redirection of lipid flux toward membrane biosynthesis via decreased ATP synthase activity, collectively causing proliferation of the ER/nuclear membrane (Han & Carman, 2017; Park et al., 2015). In *Sc*, deletion of *PAH1* improved secretion of endoglucanase CelA, β-glucosidase BglI and scFv4-4-20-mRFP (Besada-Lombana & Da Silva, 2019). Notably, *Sb*Δ*pah1* increased secretory NPA production 134%, without causing any impairment in growth (**Figure 5, S5**).

**Figure 5.**
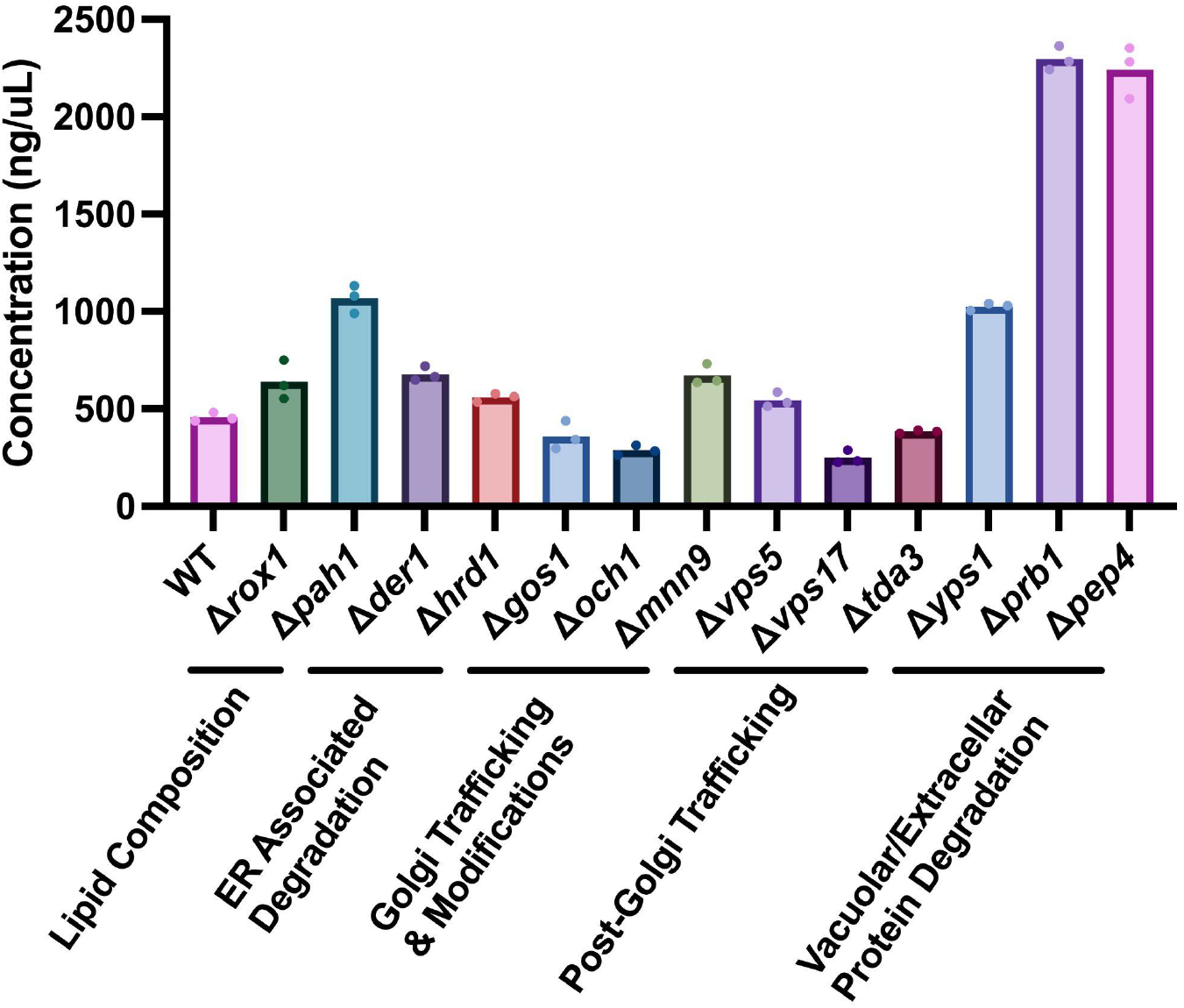
Effect of secretory pathway knockouts on peptide secretion in *Sb*. Gene candidates involved in yeast’s secretory pathway were deleted from the *Sb* genome, and the resulting strains were used to express NPA with αMF secretion signal on a high-copy (2μ) yeast origin. Each strain was cultured in triplicate in FlowerPlates in a Biolector II for 48 hours at 37 °C. Bars represent the average NPA concentration across three cultivations, and dots represent the NPA concentration in each cultivation.

**Figure S5.**
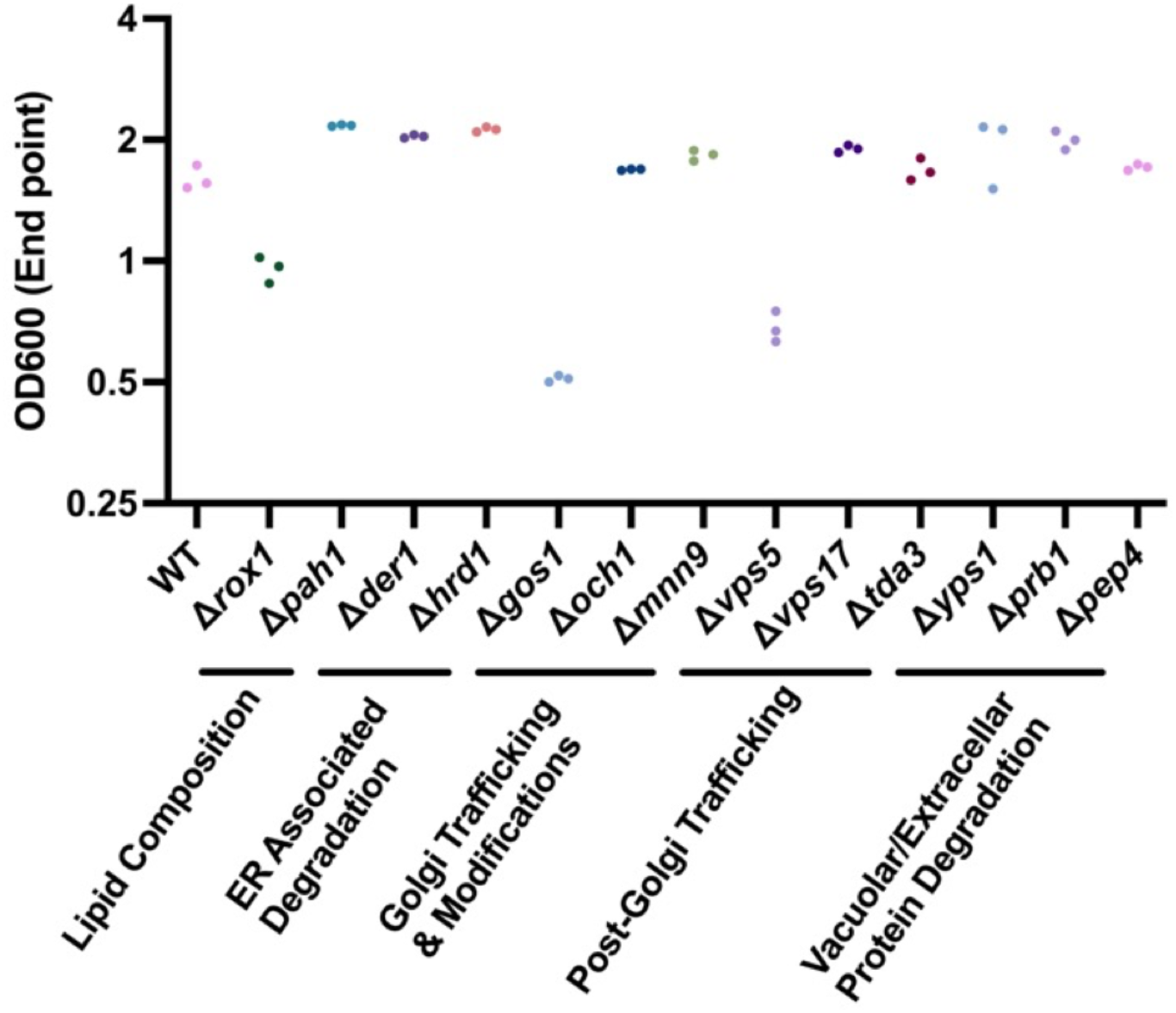
Final cell density of NPA secretor strains with varying secretory gene deletions. Each strain was cultured in triplicates in FlowerPlates in a Biolector II for 48 hours at 37 °C. Each dot represents the endpoint OD600 for each cultivation.

Second, we investigated how deletion of genes involved in ERAD processes would change NPA secretion in *Sb*. Der1p and Hrd1p are subunits of a protein complex on the ER membrane that regulates retro-translocation of aberrant proteins from the ER lumen to the cytoplasm via ubiquitination and pore formation (Igbaria et al., 2019; Knop et al., 1996; X. Wu et al., 2020). We hypothesized that impairment of ERAD through deletion of *DER1* and *HRD1* would increase NPA availability in ER and its trafficking to later stages of the secretory pathway. Deletion of *DER1* in *Sc* improved secretion of endoglucanase CelA, β-glucosidase BglI and scFv4-4-20-mRFP (Besada-Lombana & Da Silva, 2019). However, the effect of *HRD1* deletion on recombinant protein secretion has only been reported to date in *K. marxianus* (Shi et al., 2021). *Sb*Δ*der1* and *Sb*Δ*hrd1* increased secretory NPA production by 48% and 22%, respectively, and neither caused impairment in growth (**Figure 5, S5**).

Third, we focused on trafficking and modification steps that occur within the Golgi. Proteins that are processed correctly in the ER lumen are transported to the Golgi (and within the Golgi in the forward (anterograde) direction) via COPII vesicles (B. L. Tang et al., 2005). Once in the Golgi, proteins can be transported in the retrograde direction (i.e., from the late to early Golgi) via COPI vesicles (Gaynor et al., 1998). This retrograde trafficking serves to retain protein within the Golgi, facilitating modifications such as signal cleavage and glycan processing (Stanley, 2011; Zhang et al., 2001). On the other hand, retrograde trafficking is in the opposite direction of secretion, therefore potentially limiting the transfer of cargo protein to extracellular media. Furthermore, protein retention in the Golgi may lead to increased glycan formation, inhibiting transfer to the cell membrane (Kniskern et al., 1994). GOS1p is a SNARE protein regulating the anterograde trafficking within the Golgi (McNew et al., 1998). Deletion of *GOS1* improved α-amylase secretion in *Sc* (Huang et al., 2018). We hypothesized deletion of *GOS1* would also inhibit retrograde trafficking of NPA in *Sb* and increase NPA titers in the supernatant. However, *Sb*Δ*gos1* exhibited 21% less NPA production as well as significantly impaired final cell density (**Figure 5,S5**). This could potentially be due to disrupting the trafficking of materials important to vesicle formation (such as lipids) between the ER and Golgi, thereby impairing secretory performance as well as cell fitness (Bao et al., 2018; McNew et al., 1998).

When in the Golgi, proteins are modified through glycosylation and mannosylation. The extent of these modifications affects post-Golgi sorting and secretion efficiency (Smith et al., 1985). Furthermore, these glycans can interfere with the function of the secreted protein and increase their immunogenicity, particularly in mannosylation (Walsh, 2010). Och1p is an α-1,6-mannosyltransferase that initiates yeast-specific outer-chain biosynthesis of N-glycans (Lambou et al., 2010; Nakayama et al., 1992). After initial addition of α-1,6-mannan, this residue can be further elongated by addition of up to 10 α-1,6-mannan residues by Mnn9p, a mannosyltransferase (Striebeck et al., 2013). We hypothesized that deletion of these two mannosyltransferases would improve secretory production in *Sb*. Deletion of *OCH1* or *MNN9* improved secretory titers and enzyme activities of β-glucosidase, endoglucanase and cellobiohydrolase in *Sc* (*H. Tang et al., 2016*). In *Sb*, *OCH1* deletion decreased secreted NPA concentration by 37% whereas *MNN9* deletion increased secreted NPA concentration by 47% (**Figure 5**). This result shows that the same knockouts generated in closely-related strains may yield different phenotypes.

After processing in the Golgi, proteins that are correctly folded and processed are trafficked to the cell membrane to be secreted. However, some proteins are missorted to the vacuole where they are degraded (Agaphonov et al., 2005). This missorting decreases secretory titers in yeast hosts. Vacuolar missorting is regulated by several vacuole protein sorting (VPS) proteins that function as receptors directing proteins from the Golgi to early endosomes that will mature into vacuoles later on (Conibear & Stevens, 1998). Vps5p and Vps17p are subunits of a retromer complex that redirects transmembrane sorting receptor Vps10p back to the Golgi (Worby & Dixon, 2002). Vps10p is the receptor that initiates missorting to the vacuole. We hypothesized that by deleting *VPS5* and *VPS17*, we would impair vacuole missorting, as the supply of VPS10p back to the Golgi would be inhibited. In *Sc*, deletion of *VPS5* or *VPS17* improved α-amylase production (Huang et al., 2018). *Sb*Δ*vps5* exhibited 19% more secretory NPA production but had substantially impaired growth (**Figure 5, S5**). *Sb*Δ*vps17* strain exhibited 46% less secretory NPA production without a growth deficiency (**Figure 5, S5**).

We also deleted *TDA3*, which encodes for an oxidoreductase that regulates material transfer from the late endosome to the Golgi. In *Sc*, deletion of this gene improved Kex2 protease localization in the Golgi, improving the processing of pro-regions of secretion signals that contain Kex2 recognition sites such as αMF-prepro signal and consequently improving α-amylase production (Huang et al., 2018; Kanneganti et al., 2011). In *Sb*Δ*tda3*, however, secretory NPA production decreased 16% (**Figure 5**).

Fourth, we investigated knockouts of proteins involved in vacuolar end extracellular protein degradation (Hecht et al., 2014)(Ogrydziak, 1993), as these activities reduce secreted protein titers. We selected 3 proteases: Yps1p, Prb1p and Pep4p, to delete. Yps1p is an aspartic protease that cleaves proteins at mono- and paired-basic residues. It is found on the plasma membrane and in the extracellular medium. Pep4p and Prb1p are also known as vacuolar proteases A and B, respectively, and exhibit broad substrate specificities. Both are mainly present in the vacuole, but Prb1p is also present extracellularly. Deletion of these 3 proteases alone or in combination improved secretion of a human serum albumin (HSA) - human parathyroid hormone fusion protein in *Pp* and three cancer-targeting affibodies in *Sc (Gast et al., 2022; M. Wu et al., 2013*). In *Sb*, deletion of *YPS1, PRB1* and *PEP4* improved secretory NPA concentration by 124%, 402% and 390%, respectively (**Figure 5**). None of these knockouts caused growth deficiency, although the secretory production significantly increased (**Figure S5**). Vacuolar proteases are essential regulators of protein abundance and amino acid recycling in the cell, promoting cellular homeostasis (Jones, 2002). We speculate that the several additional proteases in the cell (e.g., Prc1p, Lap4p, and Cps1p) compensate for the knockouts we made, providing intracellular material and resource balance (Hecht et al., 2014).

Overall, via single-gene knockouts along various steps in the secretory pathway, we achieved between 248 and 2297 mg/L secreted NPA. It was clear that deleting the proteases enabled the highest NPA secretion, compared to the other deletions we tested (**Figure 5**), with the best-performing knockout (Δ*prb1*) improving NPA secretion 5-fold over wild-type. Crucially, several knockouts that improved protein secretion in *Sc* did not translate to *Sb*, reiterating the need for re-identifying regulators of protein secretion in this probiotic. In the future, high-throughput screening of single gene deletion or knockdown libraries will likely uncover novel engineering targets. Looking forward, these hyper-secreting strains are promising templates for expression of a wide variety of proteins of biomedical and biomanufacturing interest (Adames et al., 2019; de Ruijter et al., 2017; Huang et al., 2015; Sheng et al., 2017).

### 3.5. Combinatorial and proteomics-driven engineering improves recombinant antitoxin peptide secretion in *Sb*

We next hypothesized that knocking out several proteases simultaneously would further improve the secretory profiles we observed in single protease deficient strains. We therefore created double (Δ*pep4*Δ*prb1*) and triple (Δ*pep4*Δ*prb1*Δ*yps1*) protease knockout strains. Combining the two most beneficial protease knockouts in *Sb*Δ*pep4*Δ*prb1* improved NPA secretion by 527% without any growth restriction (**Figure 6,S6**), a 1.3- and 1.2-fold increase over *Sb*Δ*pep4* and *Sb*Δ*prb1*, respectively. However, *Sb*Δ*pep4*Δ*prb1*Δ*yps1* showed 1.7-fold lower NPA secretion than *Sb*Δ*pep4*Δ*prb1*, although the triple knockout still achieved 3.7-fold higher NPA secretion than wild-type. Studies in *Sc* have also shown that knockouts can have non-additive and difficult-to-predict effects on protein secretion (Huang et al., 2018). Therefore, we wondered whether other protease deletions may be more beneficial than *Sb*Δ*pep4*Δ*prb1*Δ*yps1*.

**Figure 6.**
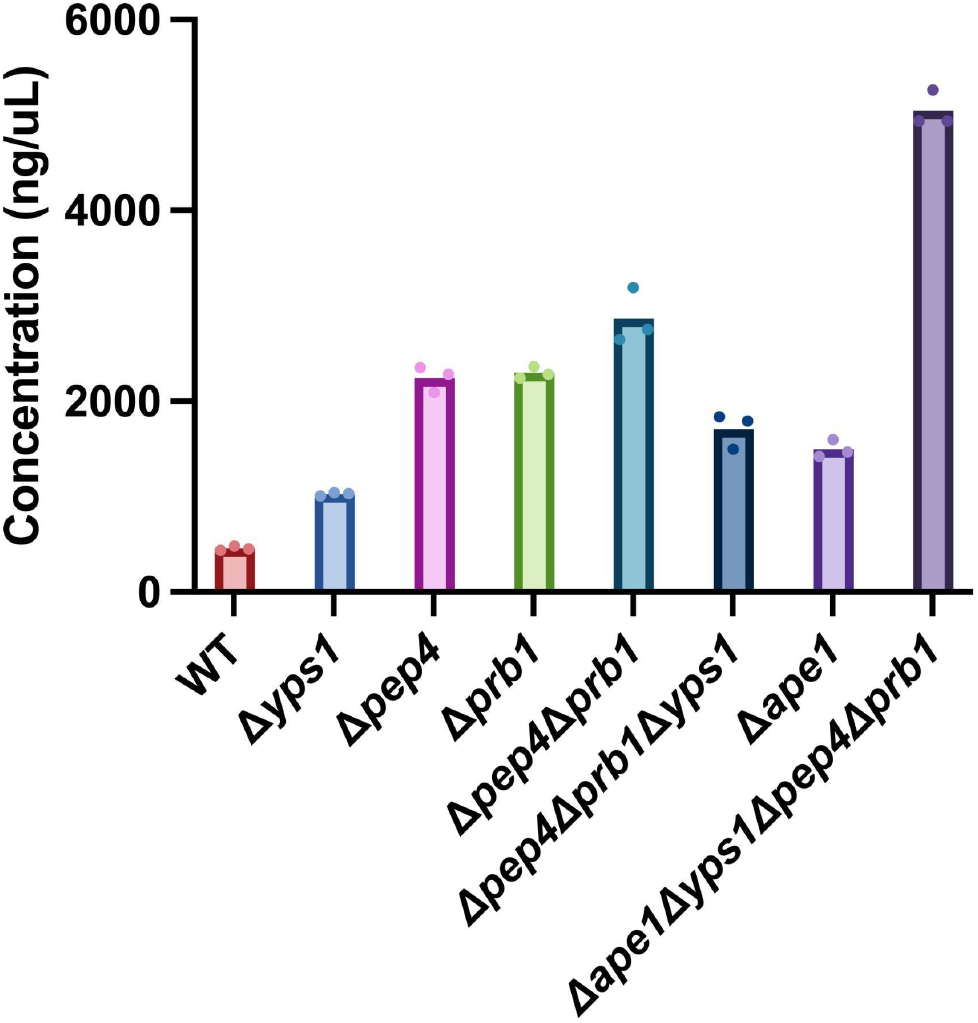
Effect of combinatorial and proteomics-driven genome engineering on peptide secretion in *Sb*. Combinatorial protease knockouts were constructed and the resulting strains were used to express NPA with αMF secretion signal from a high-copy (2μ) yeast origin. Each strain was cultured in triplicate in FlowerPlates in a Biolector II for 48 hours at 37 °C. Bars represent the average NPA concentration in three cultivations, and dots represent the NPA concentration in each cultivation.

**Figure S6.**
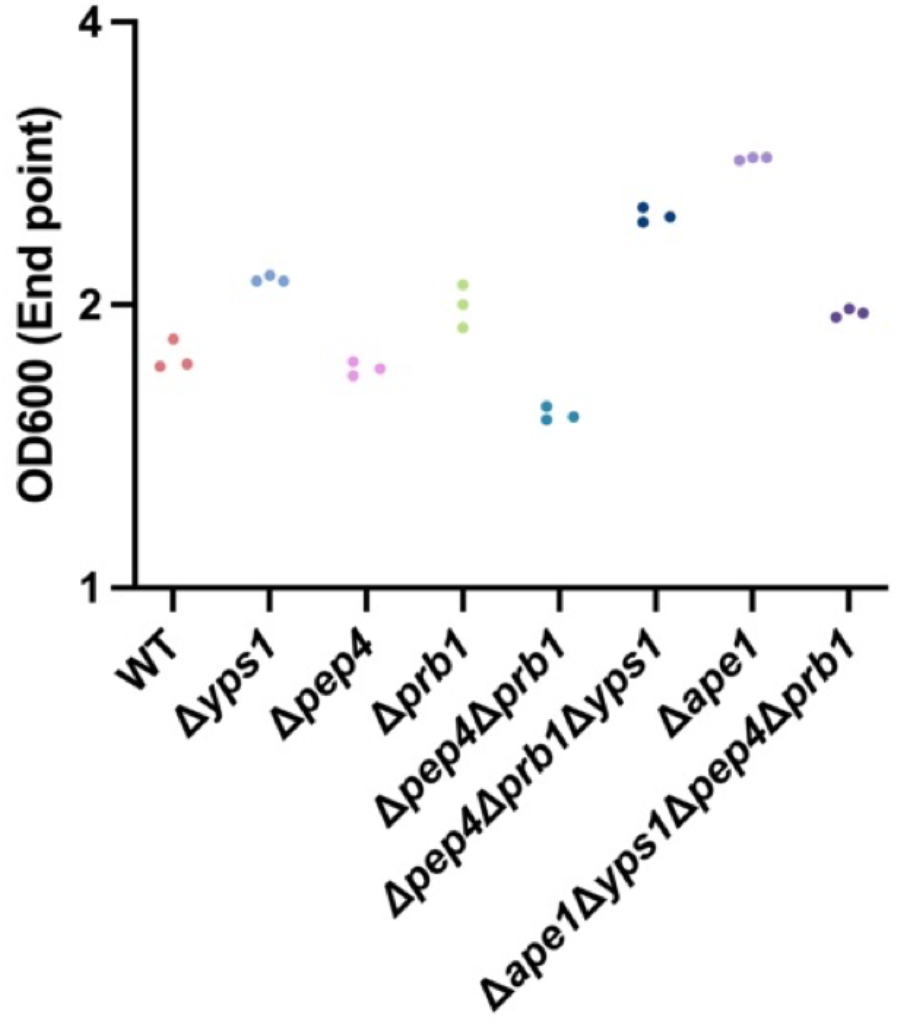
Final cell density of NPA secretor strains with combinatorial secretory gene deletions. Each strain was cultured in triplicates in FlowerPlates in a Biolector II for 48 hours at 37 °C. OD600 values were obtained at the end of the cultivation via a spectrophotometer. Dots represent the OD600 values in each cultivation for a given strain.

We hypothesized that additional proteases were produced by *Sb* (beyond Pep4p, Prb1p, and Yps1p) that limited NPA secretion. With the rationale that proteases are often present in the extracellular space, we conducted a proteomics study on supernatants collected from Sb-NPA. Precipitated supernatants were prepared for LC-MS/MS as described in the methods section. LC-MS/MS results confirmed the presence of Pep4p, Prb1p, and Yps1p in the samples (Table S5).However, we also observed a proteomic signature corresponding to Ape1p, a vacuolar aminopeptidase, that cleaves neutral or hydrophobic residues on the N-terminus (Su et al., 2015). Unlike the other 3 proteases, deletion of *APE1* has not been pursued for improvement of recombinant protein production in yeast before, to our knowledge. Deletion of *APE1* alone improved *Sb*’s secretory NPA concentration by 227%, compared to WT Sb-NPA (**Figure 6**). In addition, this deletion improved the final cell density (**Figure S6**). This could be explained by a reduced burden imposed on the cell by the cytoplasm-to-vacuole targeting (Cvt) pathway, as Ape1p is the major cargo protein for Cvt pathway (Klionsky et al., 1992; Torggler et al., 2016). When we deleted *APE1* in *Sb*Δ*pep4*Δ*prb1*Δ*yps1*, we observed substantially improved secretory NPA concentration (by 1004% compared to WT Sb-NPA), achieving NPA titers of 5045 mg/L (**Figure 6**), without exhibiting growth deficiency. (**Figure S6**).

We next investigated the effect of copy number and secretion leader on NPA secretion in the quadruple knockout strain (**Figure S7**). As observed in WT *Sb*, using a genomically integrated NPA cassette also resulted in lower NPA titers compared to an episomal vector (**Figure S7a**). However, genomically integrating NPA in the quadruple knockout strain yieIded 13.2-fold higher secretion than genomically integrating NPA in WT *Sb*. Interestingly, unlike what was observed in WT *Sb*, use of the synthetic leader preOST1-proαMF (I) did not increase the secretory production of NPA in quadruple knockout strain (**Figure S7**c), pointing to the need to tailor the secretion leader to the genotype of the chassis strain. Taken together, by harnessing combinatorial and ‘omics-driven engineering approaches, we generated an *Sb* strain with a unique genotype (a quadruple protease knockout) demonstrating significantly improved secretory performance. In future, harmonizing omics-driven and computer-guided secretory pathway models can be utilized to further identify novel combinatorial targets to improve production of proteins in *Sb* (*Huang et al., 2017; F. Li et al., 2022*).

**Figure S7.**
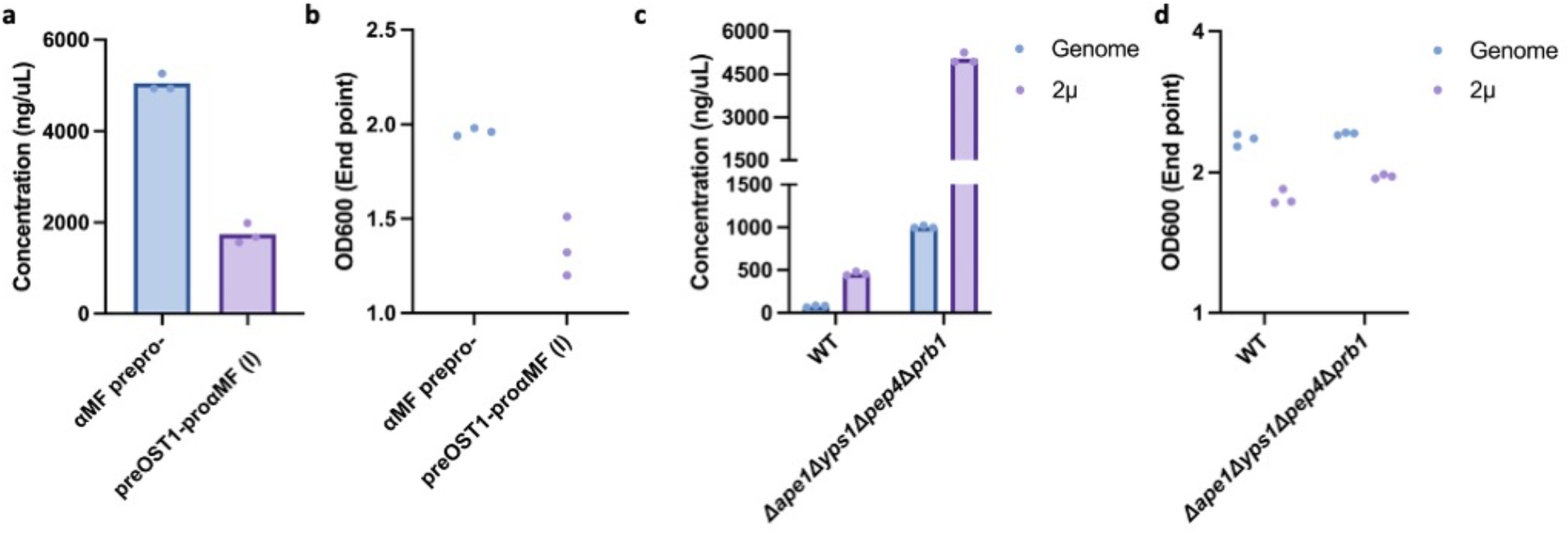
Effect of secretion signal and copy number on NPA secretion and final cell density in the quadruple knockout *Sb* strain. (a) NPA concentration in supernatant in the quadruple knockout *Sb* strain expressing NPA via the αMF secretion signal (blue) or preOST1-proαMF(I) secretion signal (purple). (b) Final cell density (OD600) for cultures in panel (a). (c) NPA concentration in culture supernatant in wild-type and quadruple knockout *Sb* strains expressing NPA via the αMF secretion signal integrated into INT1 site on the *Sb* genome (blue) or cloned into a high-copy (2*μ*) yeast plasmid (purple). (d) Final cell density (OD600) for cultures in panel (c). Each strain was cultured in triplicates in FlowerPlates in a Biolector II for 48 hours at 37 °C. OD600 values were obtained at the end of the cultivation via a spectrophotometer. For NPA concentration plots, bars represent the average NPA concentration across three cultivations and dots represent the NPA concentration in each cultivation for a given strain. For OD600 plots, dots represent the OD600 values in each cultivation for a given strain.

## 4. Conclusions

In order for therapeutic proteins to bind to human cells or other microbes in the gut, they must be secreted by the eLBP that produces them. In this work, the probiotic yeast *S. boulardii* was engineered to secrete a peptide (NPA) that targets *C.difficile* Toxin A. We first observed that encoding NPA on a plasmid vector with a high copy origin yielded 6-fold higher NPA compared to genomic integration. We next improved the secretion of NPA by exploring several native and synthetic secretion signals. Of these, preOST1-proαMF (I) enabled the highest NPA titers, increasing secreted NPA by 1.84-fold compared to the native *Sb* αMF-prepro secretion signal. Then, by individually knocking out 13 genes in *Sb*’s secretory pathway, we obtained protease knockouts that improved secretory NPA titers 5-fold over wild-type *Sb*. Finally, we harnessed a combinatorial engineering approach guided by proteomics to construct a multi-protease deficient *Sb* strain that increased secretory NPA titers 11-fold compared to wild-type. These secretion rates compare favorably to other probiotic bacteria; to our knowledge the highest protein secretion levels yet published for *L. lactis* is ~10 mg/L (Zurita-Turk et al., 2020) and 700 mg/L for *E. coli* (*Z.-Y. Chen et al., 2014; Qian et al., 2008*).

It is likely that the *in situ* performance of these engineered yeast strains will be impacted by factors such as host diet, genotype, and microbiota composition. In the future, it will be necessary to explore the relationships between the rates of *in vitro* and *in situ* biomanufacturing to facilitate rapid prototyping of therapeutic strains. We also expect that the optimal level of *in situ* secretion depends on the specific balance between on- and off-target binding kinetics for each therapeutic protein. Therefore, these results expand the range of therapeutic proteins that may be efficaciously delivered by *Sb* and open the door for the suite of high-throughput yeast engineering techniques to be used to enable further control and enhancement of therapeutic cargo delivery.

## Acknowledgements

We thank members of the Crook and Martinez-Ruiz labs for valuable discussions and input. We also thank Pablo Torres Montero and Dr. Clara Navarrete Román for help with Biolector training and experimental design. We also thank Dr. John E. Dueber for kindly sharing the MoClo-YTK Toolkit (Addgene kit # 1000000061). This work was supported by startup funds from North Carolina State University’s Chemical and Biomolecular Engineering (CBE) Department, the National Science Foundation (CBET-1934284), and the Novo Nordisk Foundation via the NCSU AIM-Bio program (NNF19SA0035474).

## Author contributions

D.D., I.S.A, and N.C. designed and conceived the study. D.D. and I.S.A. performed all yeast engineering and cultivation experiments. T.I.W. and L.B.C. performed proteomics experiments and analysis. N.C. and J.L.M.R. supervised the research. D.D., I.S.A., T.I.W., L.B.C., J.L.M.R., and N.C. wrote the manuscript.

